# Modeling mutational effects on biochemical phenotypes using convolutional neural networks: application to SARS-CoV-2

**DOI:** 10.1101/2021.01.28.428521

**Authors:** Bo Wang, Eric R. Gamazon

## Abstract

Biochemical phenotypes are major indexes for protein structure and function characterization. They are determined, at least in part, by the intrinsic physicochemical properties of amino acids and may be reflected in the protein three-dimensional structure. Modeling mutational effects on biochemical phenotypes is a critical step for understanding protein function and disease mechanism as well as enabling drug discovery. Deep Mutational Scanning (DMS) experiments have been performed on SARS-CoV-2’s spike receptor binding domain and the human ACE2 zinc-binding peptidase domain – both central players in viral infection and evolution and antibody evasion - quantifying how mutations impact binding affinity and protein expression. Here, we modeled biochemical phenotypes from massively parallel assays, using convolutional neural networks trained on protein sequence mutations in the virus and human host. We found that neural networks are significantly predictive of binding affinity, protein expression, and antibody escape, learning complex interactions and higher-order features that are difficult to capture with conventional methods from structural biology. Integrating the intrinsic physicochemical properties of amino acids, including hydrophobicity, solvent-accessible surface area, and long-range non-bonded energy per atom, significantly improved prediction (empirical p<0.01) though there was such a strong dependence on the sequence data alone to yield reasonably good prediction. We observed concordance of the DMS data and our neural network predictions with an independent study on intermolecular interactions from molecular dynamics (multiple 500 ns or 1 μs all-atom) simulations of the spike protein-ACE2 interface, with critical implications for the use of deep learning to dissect molecular mechanisms. The mutation- or genetically-determined component of a biochemical phenotype estimated from the neural networks has improved causal inference properties relative to the original phenotype and can facilitate crucial insights into disease pathophysiology and therapeutic design.

## Introduction

Since the initial outbreak, the SARS-CoV-2 virus has rapidly spread worldwide causing a global public health crisis, the coronavirus disease 2019 (COVID-19). *In vitro* and cryo-electron microscopy studies have established that the betacoronavirus uses the human cell-surface protein angiotensin converting enzyme 2 (ACE2) to gain entry into target cells^1–3^. Therefore, precise characterization of the interaction between the Receptor Binding Domain (RBD) of the viral spike glycoprotein and the ACE2 complex is of critical importance in understanding COVID-19 pathophysiology^3^. Not surprisingly, several drug candidates that target either the virus or the receptor have been developed on the basis of the ACE2 binding. With improved understanding of this key molecular interaction, two major therapeutic strategies have been pursued, including 1) engineering high-affinity ACE2 decoy or developing antibody cocktail treatments and 2) screening new or repurposing existing inhibitors targeting the binding interface^4,5^. Establishing the sequence-structure-phenotype relationship for the spike RBD and the ACE2 receptor is essential for both strategies, in which the sequence mutational effect on receptor affinity and other biochemical phenotypes is the major component^6–10^.

Comprehensive understanding of how variants, including single mutations, affect disease-relevant biochemical phenotypes would go a long way towards clarifying molecular mechanisms of disease as well as downstream adverse complications and guiding pharmacological interventions. In addition, elucidating the mutational effect may shed light on selective pressures determining the evolutionary trajectory of the coronavirus as well as identify risk factors for viral infection and maladaptive host response to COVID-19 in human populations^11^. Deep Mutational Scanning (DMS) systematically evaluates the effect of mutant versions of the protein on measured biochemical phenotypes ^12,13,6,14^. High-throughput mutagenesis in DMS makes it possible to assess the phenotypic consequences of each possible amino acid mutation in a protein, generating large datasets that can reveal the sequence-function landscape. The development of computational approaches to learn the complex and non-linear features of this map can enable high-throughput inference of basic protein properties. Statistical and machine learning methods, including deep learning, have attracted significant attention due to their predictive power^15^. A recently developed supervised learning framework tailored to DMS datasets, convolutional neural networks demonstrated spectacular performance, consistent with other recent studies of mutational effect^16,17^.

DMS experiments on both the SARS-CoV-2 spike glycoprotein and the ACE2 receptor have been performed, providing an important basis for further investigations of mutational effects ^4,7,18^. In this work, we conducted systematic modeling of the mutational effects of the RBD in the viral spike protein and of the ACE2 receptor on biochemical phenotypes, extending a supervised learning framework^16^. Three classes of critical phenotypes -- binding affinity, protein expression, and antibody escape -- were systematically analyzed within the sequence-structure-function paradigm that informs much of proteomic and structural biology studies. Neural networks were also leveraged to learn (a) the complex functional landscape of the viral spike protein’s RBD and the host cell-surface receptor ACE2 and (b) the antibody-escape map of RBD mutations, including mutations that undergo selection during viral proliferation in the presence of antibodies and mutations that have been circulating in human populations^19–21^. Finally, the mutation- or genetically-determined phenotypes (i.e., the component of a phenotype defined by the sequence versus a technical confounder or the environmental component) estimated from the neural networks were exploited to extract key insights into the molecular mechanisms of COVID-19 infection and severity.

## Results

### Overview of framework

Briefly, we describe the framework (Figure 1A). Here we consider a set of N training samples consisting of (protein) sequences, x_1_, x_2_, …, x_N_ ∈ X, with corresponding biochemical phenotype values, y_1_, y_2_, …, y_N_ ∈ Y (where phenotype is fixed and chosen from binding affinity, protein expression, or an antibody-escape measure), where the pairs are sampled independently and identically distributed (i.i.d.) from a joint distribution

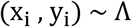

**Figure 1.**
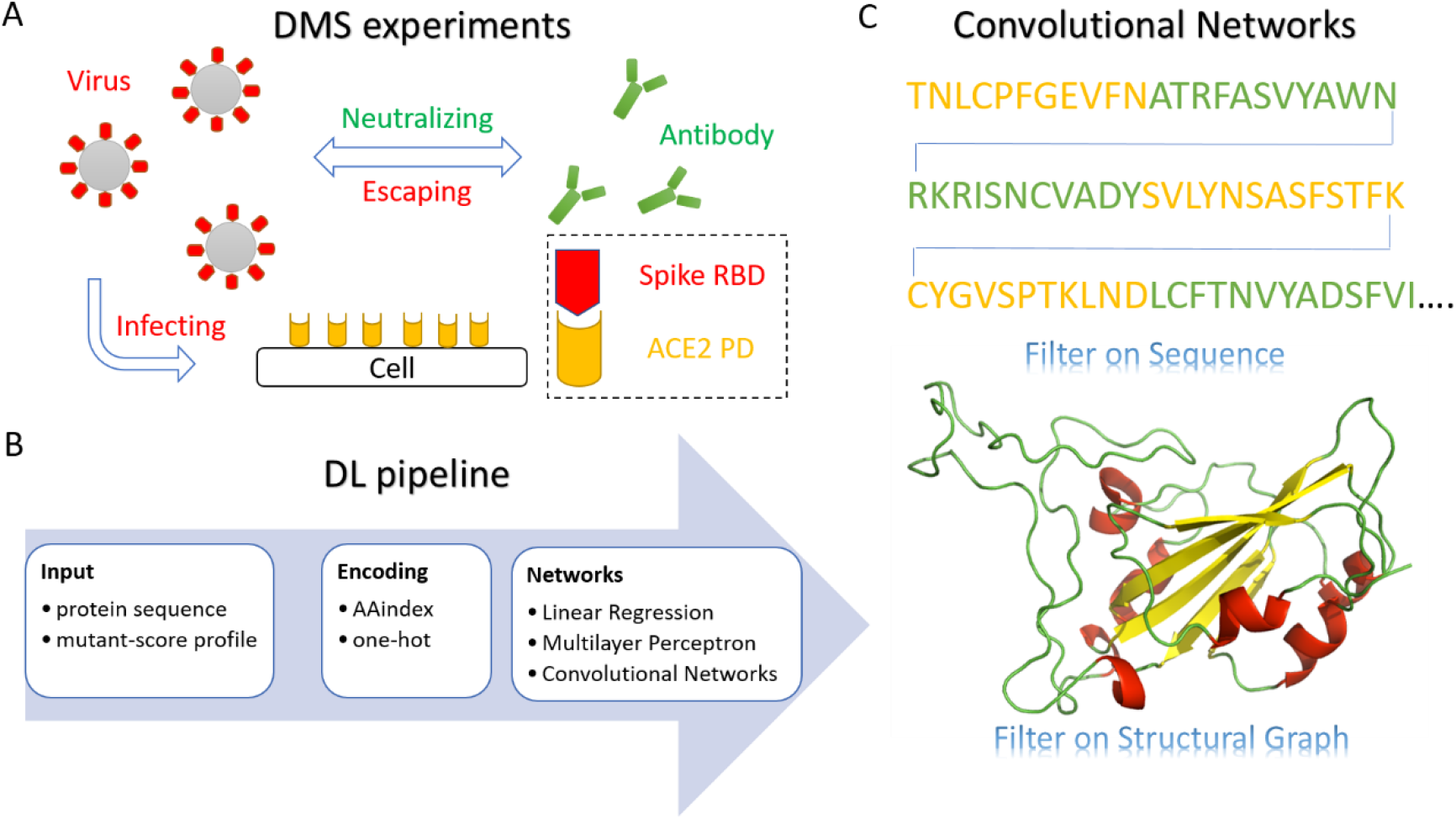
Overview of the framework. **A** We evaluated three biochemical phenotypes generated from Deep Mutational Scanning (DMS) experiments: binding affinity, protein expression, and antibody escape. These biochemical phenotypes may determine the evolutionary trajectory of the coronavirus, viral infection, and maladaptive host response. **B** We quantified the extent to which state-of-the-art neural network methodologies can predict these phenotypes, using only sequence mutation data. The predicted biomedical phenotypes are mutation- or genetically-determined traits, which are therefore of intrinsic interest because they are determined solely by the (amino-acid) sequence data. We encoded the protein sequence input using AAindex, consisting of 566 intrinsic (data-independent) physicochemical properties of the amino acids, such as hydrophobicity and long-range non-bonded energy per atom, and one-hot encoding. We then trained neural networks on the mutation-phenotype data from the DMS experiments. **C** We trained sequence convolutional neural networks and graph convolutional neural networks (along with [fully connected] multilayer perceptrons and linear regression) on the DMS data. Shown are the convoluted sequences and structural motifs for the two types of architecture, respectively, differentiated by color.

A supervised learning method seeks to find, from a pre-defined class of functions, a function g: X →Y, which maps the input sequence × to the phenotype value *y* and minimizes the expected deviation

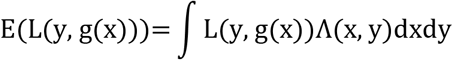

from *y* under a loss function L: Y × Y → R. Regularization (e.g., dropout) is also performed to prevent overfitting and enhance generalization error. The expected loss is generally approximated by the average of the L(y_*i*_, g(x_*i*_)) across the training samples, i.e., the empirical loss, given that the distribution Λ is typically not known. Neural networks constitute the class of functions defined by the composition of affine maps (Ψ_i_) and activation functions (φ) (which are typically non-linear).

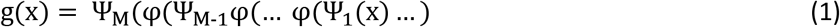

(Note linear regression can be trivially included as a member of this class of functions.) The minimization problem, i.e., the search for an optimal function g, is framed as a search for a set of weights ***W***(_*k*_) and biases *b*(_*k*_) (Methods).

### Application of deep learning framework to the spike RBD and the ACE2 PD complex

The crystal structure of the spike RBD — ACE2 PD complex was retrieved from the Protein Data Bank (PDB; PDB ID: 6M0J)^2^. Our study investigated two biochemical phenotypes, binding affinity and protein expression – each critical to viral evolution, receptor recognition, and host-virus interaction – and a set of antibody-escape phenotypes, which may provide critical insights into the antigenic consequences of mutations and effective vaccine design (Figure 1A). Specifically, we trained on single-mutation functional score profiles, consisting of all possible amino acid mutations and the corresponding phenotypic measurements (quantified relative binding score for the ACE2 catalytic zinc-binding peptidase domain (PD), binding affinity and expression for the spike RBD, and the “escape fraction” from each of 10 human monoclonal antibodies) measured by DMS experiments. The protein sequences x_i_ were encoded by one-hot encoding and an external curated featurization table, AAindex, which summarizes the intrinsic physicochemical properties of amino acids, e.g., hydrophobicity, solvent-accessible surface area, and long-range non-bonded energy per atom (Figure 1B; Methods).

We generated independent models, which allow prediction of mutational effects on binding affinity, expression, or antibody escape (for each of the 10 antibodies) of previously uncharacterized variants. We implemented linear regression and several neural network architectures: multilayer perceptron (fully connected), sequence convolutional neural network, and graph convolutional neural network. While linear regression considers only the additive contributions of sequence residues, multilayer perceptron can capture nonlinear relationships between sequence (input) and phenotype (output). Due to their distinctive architecture, convolutional neural networks are even more powerful to learn higher-level features. Filters for sequence convolutional neural networks slide along the one-dimensional input of protein sequence and learn from sequence fragments while graph convolutional neural networks act on the structural coordinates of proteins (Figure 1C). In the graph structure representation G = (V, E) of the spatial structure of the protein, the sequence residues are represented as nodes *V* while the interactions for pairs of residues (determined by a distance threshold) are represented as edges *E*. Graph convolutional neural networks learn segments of protein structures, i.e., motifs, with the convolutional kernels capturing translational invariance.

### Evaluating models learned from Deep Mutational Scanning data

We performed the supervised learning methods on the various spike RBD and ACE2 PD datasets, comprising mutant-score profiles of binding affinity, expression, and antibody escape. For each model (defined by a phenotype from a DMS dataset), the dataset was randomly split into 60% training, 20% tuning, and 20% testing subsets. Trained models were evaluated by predictive performance on the test subset (e.g., Figure 2A). The supervised learning methods demonstrated varied predictive performance in the case of the spike RBD, where the Spearman correlation coefficient between the observed and predicted phenotype ranged from 0.49 (for linear regression) to 0.78 (for convolutional neural network) for ACE2 binding affinity (Table S1). Neural networks provided much better predictive performance than linear regression, indicating that binding affinity is characterized by interactions (non-additive effects) among residues and a non-linear mapping of sequence to function. Relative to the distribution of binding affinity (Figure 2A) from the RBD mutations, the predicted phenotypes from the neural networks had greater fidelity to the original than linear regression (Figure 2B and 2C). Notably, the amino acid sequence mutational data alone could significantly predict binding affinity (p < 2.2×10^−16^), using multilayer perceptron (Spearman correlation *ρ* = 0.52), sequence convolutional neural network (*ρ* = 0.69), and graph convolutional neural network (*ρ* = 0.70) (Figure 2D, 2E, and 2F, Table S1).

**Figure 2.**
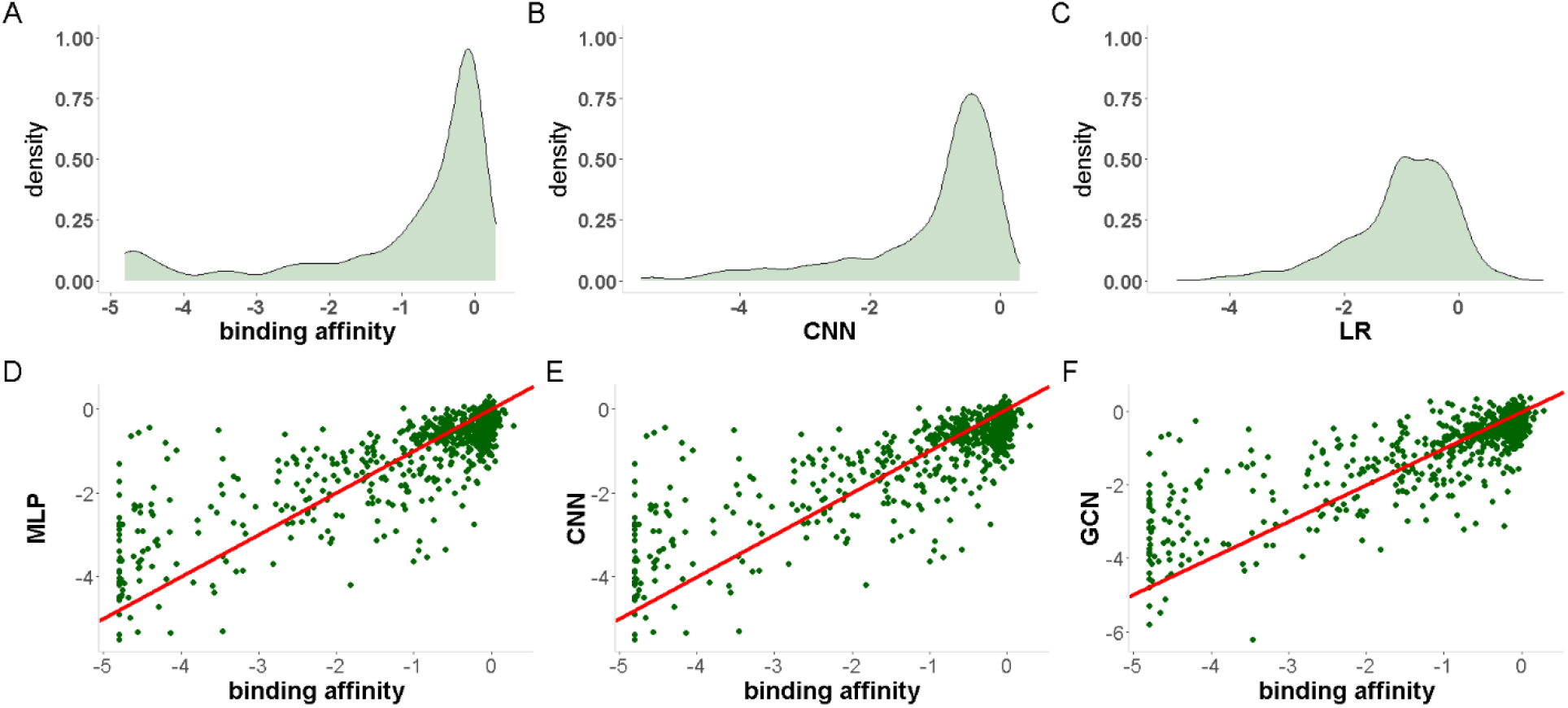
Predictive performance of neural networks on binding affinity. **A** Distribution of ACE2 binding affinity, in the test set, from single mutations in the spike glycoprotein RBD, as measured by DMS. The predicted biochemical phenotype from **B** neural networks, e.g., CNN, was much closer in distribution to the original phenotype than that from **C** linear regression. Notably, **D-F** the 3 neural network models trained here showed significant Spearman correlation (p < 2.2×10^−16^) between the observed (x-axis) and predicted (y-axis) phenotype in the (independent) test set. Red line is the diagonal (*y* = *x*) line. CNN: convolutional neural network (Table S1, Network 2); LR: linear regression; MLP: multilayer perceptron; GCN: graph convolutional network (Table S1, Network 8).

For the spike glycoprotein expression, the prediction performance (Spearman correlation in the test set) ranged from 0.44 (for linear regression) to 0.84 (for graph convolutional network). Binding affinity to ACE2 and spike glycoprotein expression were significantly correlated (*ρ* = 0.65) in the training set, but the relationship was found to be highly nonlinear (Figure 3A). Though both phenotypes reflect protein stability under a wide range of biochemical contexts, binding affinity is more determined by the surface residues on or near the binding sites as well as by allosteric effects induced by inner or remote residues. As in the case of binding affinity, the neural network models significantly predicted RBD expression (p < 2.2×10^−16^), using multilayer perceptron (Spearman correlation *ρ* = 0.50), sequence convolutional neural network (*ρ* = 0.682), and graph convolutional neural network (*ρ* = 0.679) (Figure 3B, 3C, and 3D, Table S1). Despite the range in prediction performance of the models, the estimates of the mutation-mediated component of RBD expression derived from them were highly correlated (Figure 3E).

**Figure 3.**
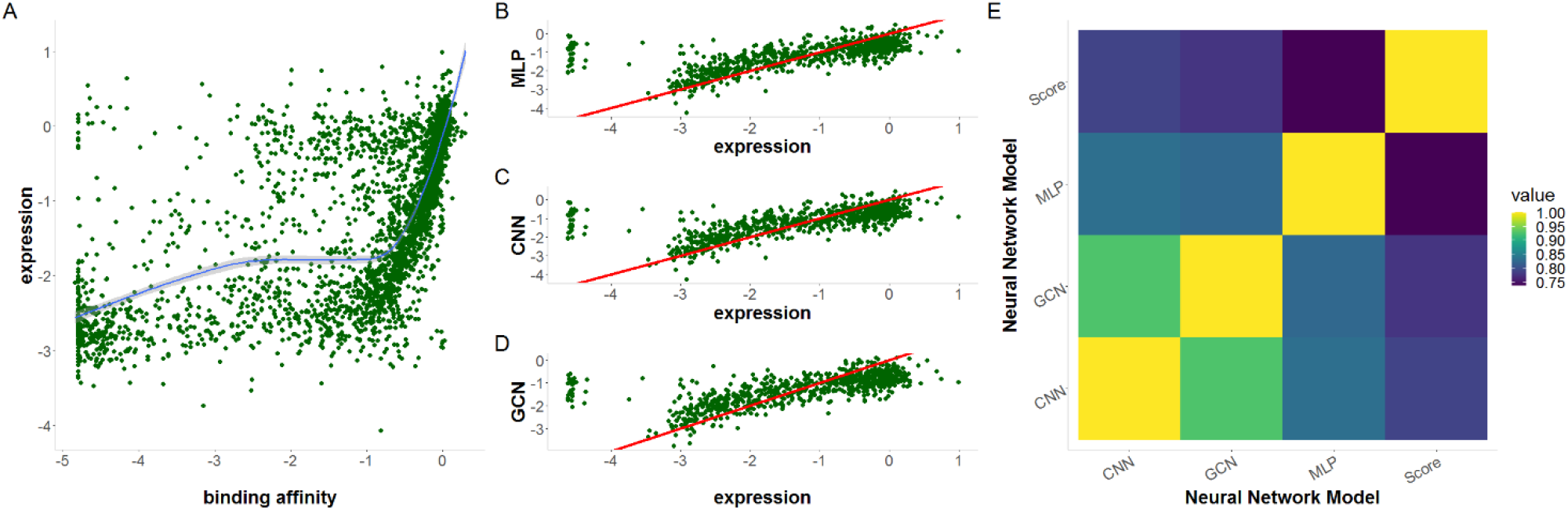
Predictive performance of neural networks on protein expression. **A** Binding affinity to ACE2 and spike glycoprotein expression, both induced by a protein sequence mutation on the RBD, were significantly correlated (Spearman correlation *ρ* = 0.65), but the relationship between the two phenotypes was highly nonlinear. We fit a curve through the points, using Locally Estimated Scatterplot Smoothing (LOESS), a method for local polynomial regression. **B-D** We trained three neural network models, which could significantly predict (Spearman correlation p < 2.2×10^−16^) protein expression in the (independent) test set. Red line is the diagonal (*y* = *x*) line. **E** Pairwise correlation between the neural network models. MLP: multilayer perceptron; CNN: convolutional neural network (Table S1, Network 2); GCN: graph convolutional network (Table S1, Network 8).

Previous studies have shown that mechanistic insight of protein mutational effects is not easily transferable to studying the effects of variants in other proteins^14^. We hypothesized that models trained on one species-phenotype are not suitable for predicting another species-phenotype. To evaluate this hypothesis, we trained a model using the avGFP DMS dataset^22^, in which the score of the green fluorescent protein from *Aequorea victoria*, avGFP, is fluorescence. The avGFP sequence was slightly modified to match the input dimension of the spike RBD (Methods). Therefore, the expression-based score profile of the spike RBD could be used as an external test set, with its combination of amino-acid sequence unseen during the training and an unrelated phenotype. As hypothesized, the sequence convolutional model from avGFP failed to predict the expression of mutated sequences (Spearman correlation *ρ* = −0.0346). This finding implies that the biochemical phenotypes differed in their dependence on protein amino-acid sequence length, type, or composition, with interaction from non-sequence physicochemical environments (for example, pH condition and ionic density) potentially accounting for the difference.

### Influence of network architectures on biochemical phenotype prediction

All mutational sequences were encoded by the externally defined AAindex, a set of 566 quantitative indices of physicochemical properties of amino acids, which were used to train the empirical mutational scores on the DMS data^23^. Thus, we examined the impact of the quality of the AAindex on prediction performance. We shuffled the entire AAindex so that the intrinsic information for each amino acid would be completely lost and replaced with random noise (Methods). We generated 100 pseudo AAindex tables, on which sequence neural networks were trained, using a given choice of configurations, i.e., number of layers and filter dimensions (Figure 4A). For each choice of network configurations, the distribution followed a bell-shaped curve, with the prediction performance from the actual AAindex located at the far-right and outperforming any of the shuffled tables. In other words, integrating the AAindex information significantly improved (empirical p<0.01) the overall performance of the neural networks.

**Figure 4.**
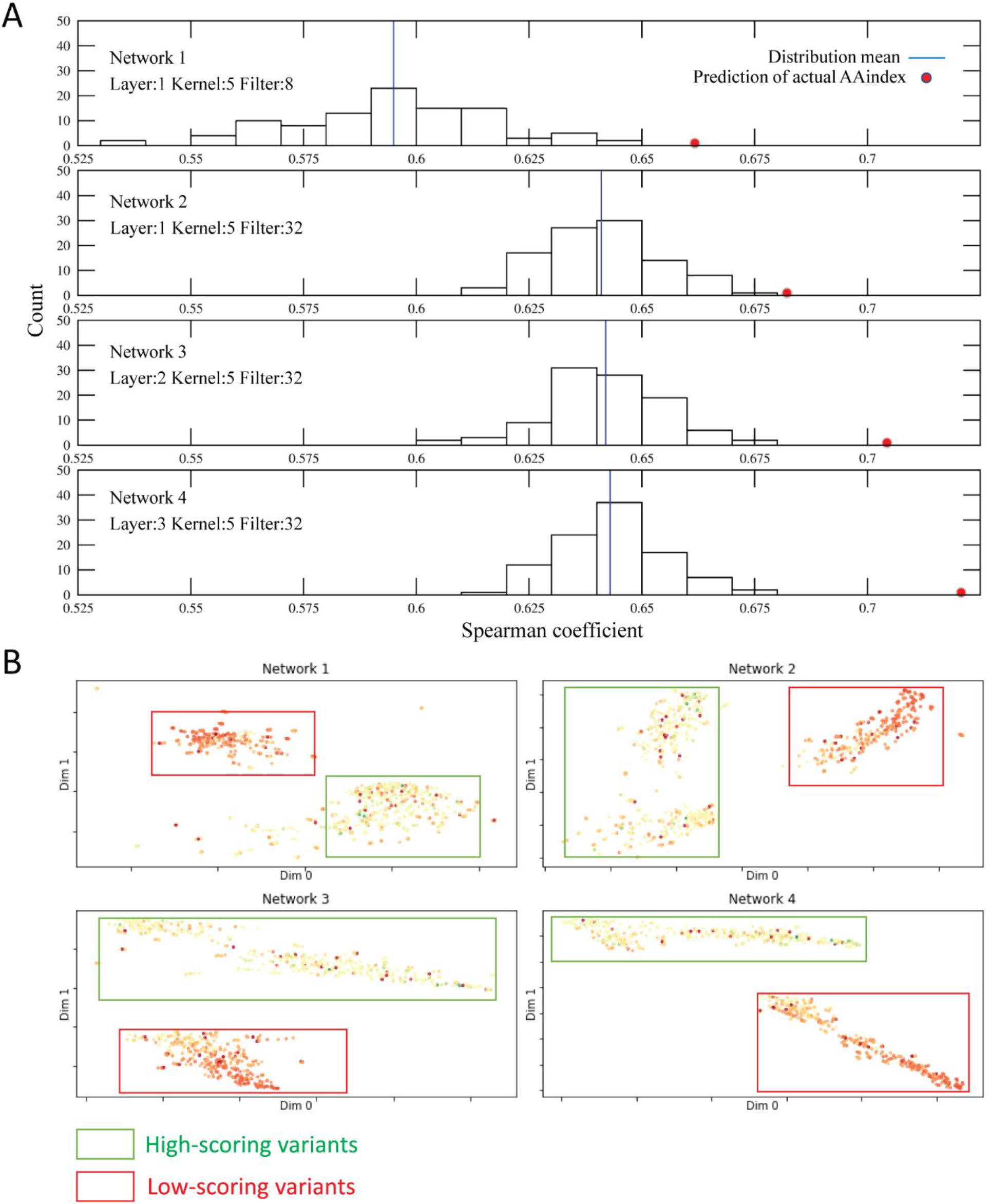
Effect of neural network architectures on predicting biochemical phenotypes. **A** Distribution of predictive performance (Spearman correlation in the test set) of sequence convolutional neural networks with a shuffled AAindex (shown as a histogram) and the actual AAindex (shown as a red dot), training on the spike RBD expression dataset. Each plot corresponds to a specific choice of architecture, including the number of layers and the filter dimension. Increasing the filter dimension or the number of layers improved the prediction performance. Furthermore, encoding the input data with the actual AAIndex, which summarizes the externally-defined intrinsic physicochemical properties of the amino acids, consistently led to performance gain. Nevertheless, we observed a strong dependence, on the input sequence data alone (i.e., without the intrinsic amino acid attributes), of the original biochemical phenotype, with reasonable performance across all choices of network configurations. **B** Uniform Manifold Approximation and Projection (UMAP) projection of the latent space of the sequence convolutional networks on the last internal layer of the network. Mutants are colored according to the actual phenotype score, with green and red representing high-scoring and low-scoring variants, respectively.

The prediction substantially improved with a larger filter dimension on a single hidden layer sequence convolutional network. This observation implies that the neural networks were able to learn more information from longer sequence fragments, which is consistent with the expression of the spike RBD reflecting complex non-linear relationships of the sequence (Network 1, 2 Figure 4A). Even with a shuffled AAindex table, i.e., in which the intrinsic properties of the amino acids were essentially lost, a larger filter dimension led to enhanced learning from the expression phenotype, with increased mean and reduced standard deviation for the empirical performance distribution, indicating highly strong dependence of expression on the sequence data. Deeper neural networks with a fixed filter dimension also showed improved prediction on expression (Network 2-4, Figure 4A). Integration of an accurate AAindex as input of a deeper neural network resulted in further performance gain relative to a shuffled AAindex, confirming that the neural network was leveraging the externally-defined intrinsic properties. To further reveal the information learned from neural networks by increasing the number of hidden layers, we applied Uniform Manifold Approximation and Projection (UMAP) to visualize the latent space in the last layer^16,24^. Notably, a deeper network could better differentiate high- and low-scoring sequence mutants as distinct clusters (Network 1-4, Figure 4B).

### Mutational effects on biochemical phenotypes

SARS-CoV-2 infection leads to complex pathophysiological consequences which are far from being fully understood. Elucidating mutational effects on biochemical phenotypes for the spike RBD and the ACE2 PD is critical in improving our understanding of the virus-host interactions as well as host immune response. Furthermore, with the spike glycoprotein as the primary target due to its role in mediating coronavirus entry into host cells, neutralizing Antibodies (Abs) may provide protection from SARS-CoV-2 in humans^25^. Here, we highlight the results from sequence convolutional neural networks that showed the optimal performance among all tested models and further interpret them in connection with existing experimental characterizations, molecular modeling, and host genetic inferences. In addition, we leveraged the neural network derived phenotypes to evaluate their utility in downstream analyses because, as inferred phenotypes, they are solely determined by the sequence mutation data and are thus less biased by possible confounding artifacts^26,27^, including batch effects, environmental (non-sequence physicochemical) determinants, and other (hidden) technical sources of heterogeneity.

#### Key mutations on spike RBD

SARS-CoV-2 evolution promotes virulence and enhances escape from neutralizing Abs. Interface mutations (Figure 5A and 5D), specifically those at contact sites, are more likely to have direct impact on infectivity. The mutational effects of these “key binding residues” of the spike RBD, i.e., residues that are essential for ACE2 binding based on structural characterization^2^, were systematically evaluated by our neural networks (Figure 5B and 5E). Predicted phenotypic effects were consistent with DMS measurements^7^; for example, mutations at G502 showed the most extreme impact on receptor binding affinity (see Figure 5B). The neural network estimate of mutation-mediated binding affinity with the ACE2 receptor significantly differed (Mann-Whitney U test p=5.19×10^−18^) between the set of key binding residues and the remaining residues in the spike RBD, thus inducing clustering. Notably, the difference between the two residue classes was more significant (p=6.9×10^−14^) for the neural network derived phenotype than for the original phenotype, i.e., the measurement unadjusted for non-mutational effects and (hidden or unmeasured) technical confounders. Interestingly, across the key binding residues in the RBD, a mutation to a Tyrosine (Y) showed higher neural network derived receptor binding affinity and lower variability compared with a mutation to another amino acid (Figure 5F). The outlier Tyrosine mutational effect on the level of receptor affinity was not detectable with the original (unadjusted) phenotype. In general, there was much higher variability across the key binding residues in the original phenotype than in the neural network estimate of the mutation-mediated phenotype (Figure 5F). In addition, the key spike RBD binding residues were consistent with results of interfacial interactions of the spike RBD bound to ACE2 from molecular dynamics simulations^28^ (performed for 500 nanoseconds in triplicates or 1 μs all-atom, using Desmond). Intermolecular interactions, including hydrophobic interactions, hydrogen bonds, π-π, and cation-π, were dynamically observed in formation and breakage in the simulations. Notably, the key binding residues of the spike RBD formed consistent polar interactions with the corresponding residues of ACE2 during the molecular dynamics simulation trajectories (Figure 5C).

**Figure 5.**
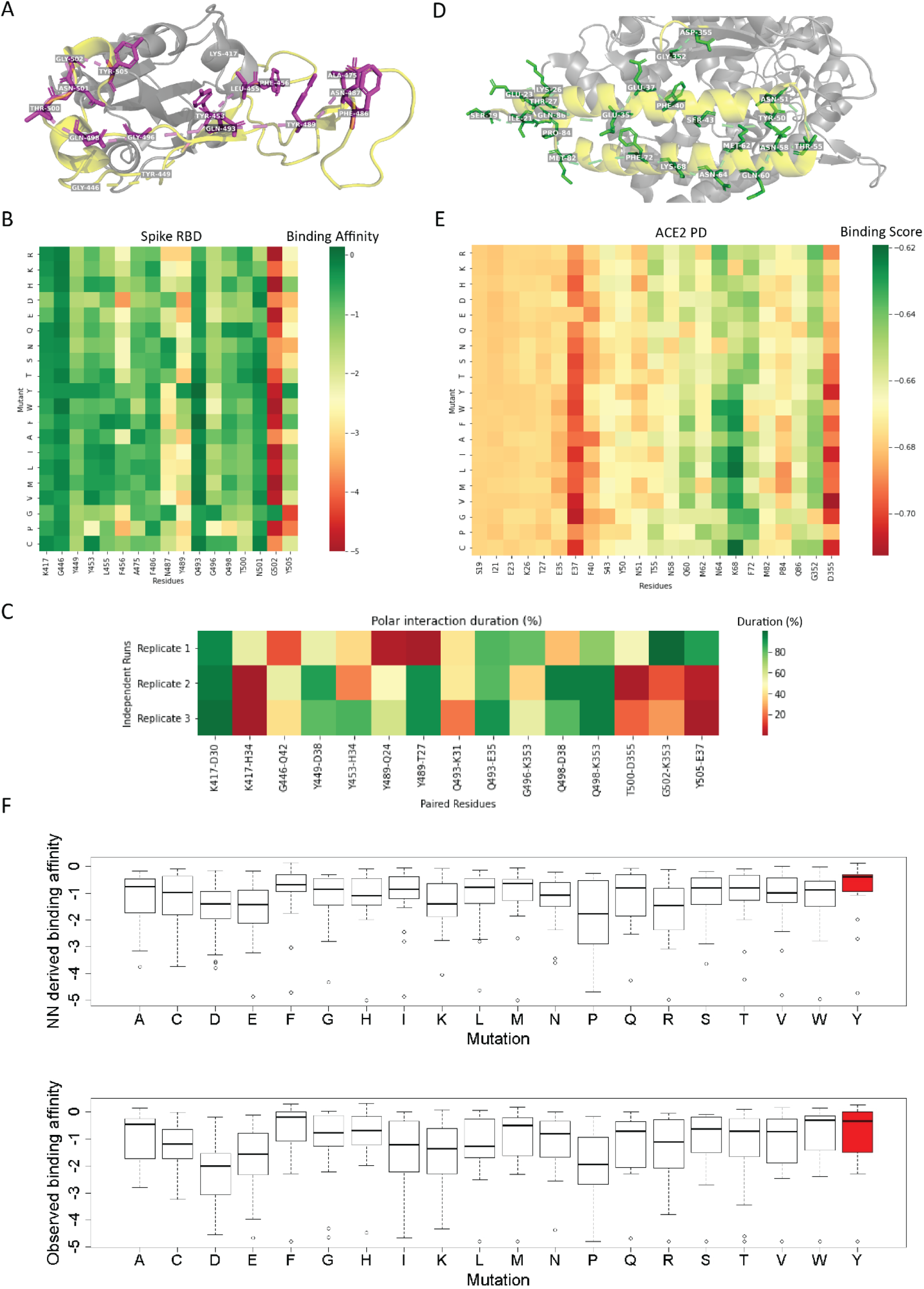
Mutational effects at key interface residues on the spike RBD and ACE2 PD. **A** Human ACE2-interacting residues (magenta) of the spike RBD interface (yellow) with **B** their predicted mutational effects on receptor affinity from a convolutional neural network with optimal performance (Network 3, Table S1). These key binding residues significantly differed (Mann-Whitney U test p=5.19×10^−18^) in the neural network derived ACE2 binding affinity from the complement set of residues in the spike RBD, with the difference between the two residue classes more significant (p=6.9×10^−14^) for this refined phenotype than for the original phenotype, i.e., the measurement unadjusted for non-mutational effects and technical confounders. Note the residues with outlier predicted mutational effects on binding affinity. **C** Paired residues between the spike RBD and the ACE2 PD that retained polar interactions and the interaction duration percentage from molecular dynamics simulation trajectories. Briefly, 500 nanosecond molecular dynamics simulations had been performed in triplicates using Desmond (ref 28), with the structure of the spike RBD bound to ACE2 solvated with single point charge water molecules and with use of the OPLS forcefield. Intermolecular interactions (e.g., hydrophobic interactions, hydrogen bonds, π- π, cation-π) were dynamically observed in formation and breakage. We leveraged the residues of the spike RBD forming consistent polar interactions with the corresponding residues of ACE2 to evaluate concordance with the DMS measurements and our neural network predictions. Each paired residues included here were in polar interaction for at least 40% of the simulation duration in at least one replicate. **D** Human polymorphisms (shown in green stick representation), based on the gnomAD resource (v2) from large-scale sequencing projects, of the ACE2 PD interface with **E** the corresponding residues’ predicted mutational effects on binding affinity from a convolutional neural network with optimal performance (Network 5, Table S2). Note the residues with outlier predicted mutational effects on relative binding score. **F** Among the key binding residues, a mutation to a Tyrosine (red) showed higher neural network derived ACE2 receptor binding affinity and lower variability compared with a mutation to another amino acid. For comparison, a similar plot for the original phenotype, unadjusted for non-mutational effects and technical confounders, is shown in the bottom panel. The outlier Tyrosine mutational effect was not detectable with the original phenotype. Furthermore, we observed, in general, higher variability across the key binding residues in the original phenotype than in the neural network estimate of the mutation-mediated phenotype.

Neutralizing Abs secreted from immune cells may also be found to be in complex with the spike RBD. Consistent with the experimental characterizations^19^, the neural network derived antibody-escape phenotypes for the 10 human monoclonal Abs have differentiated target “hotspots” on the surface of the spike RBD, pointing to different sets of escape mutations (Figure 6A). The neural network prediction performance on antibody escape of the spike RBD varied between the Abs (Spearman correlation coefficient in the test set between 0.09 and 0.32) (Figure 6B). The variability in performance might be due, at least in part, to the unique physicochemical properties of the 10 monoclonal antibodies as well as their specific relative dependence on sequence mutation and other (non-sequence) determinants.

**Figure 6.**
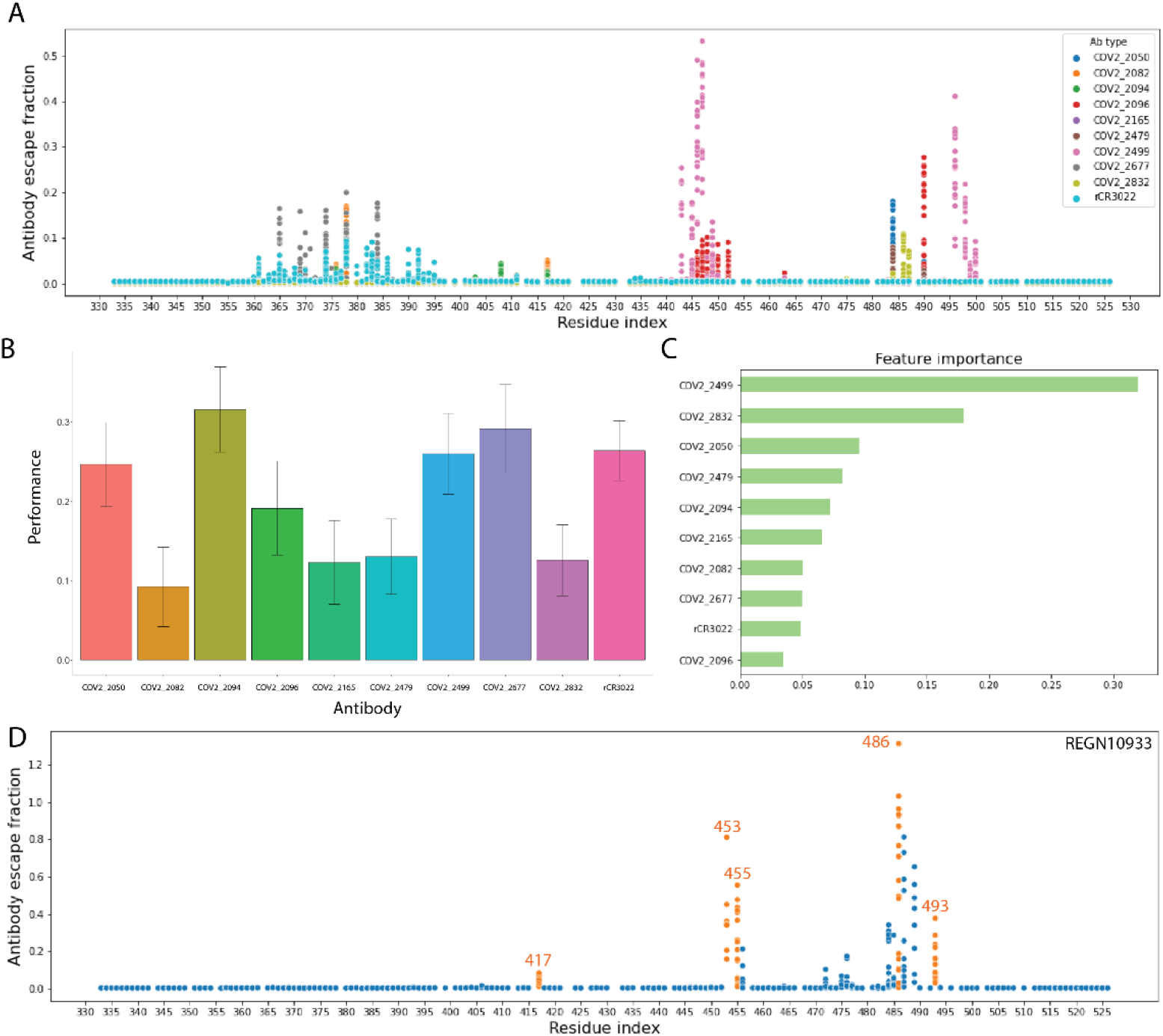
Neural network derived antibody-escape phenotypes, performance prediction, and feature importance. **A** Maps of the convolutional neural network derived (i.e., estimated, mutation-mediated) escape fraction phenotype on the spike RBD for the different Abs (shown as different colors). Shifting peaks in different colors indicate differentiated binding sites and highlight different sets of escape mutations for the Abs on the spike RBD surface. Key residues on the spike RBD that are critical binding sites with ACE2 PD include G446, Y449, G496, Q498, and T500 for escape mutations of COV2-2499. Note these residues are the sites of the peaks of the neural network derived antibody-escape phenotype for COV2-2499, i.e., the predicted escape mutations. The importance of these residues for viral binding to the host receptor is also supported by results on intermolecular interactions from the molecular dynamics simulations (Figure 5C). **B** Predictive performance (Spearman correlation in the test set between the observed and predicted antibody-escape fraction) of our neural network model on each of the 10 Abs (Table S3). For each antibody, the standard error is shown, as generated from application of *bootstrap* (n=100). **C** Feature importance of the convolutional neural network derived (i.e., estimated, mutation-mediated) antibody-escape phenotype for each of the 10 Abs as derived from a joint Random Forest regression model in predicting binding affinity towards the ACE2 receptor. The Abs that shield the receptor by entirely targeting the binding interface ranked in the top 3 neural network derived antibody-escape predictors. **D** Map of neural network derived (i.e., estimated, mutation-mediated) antibody-escape phenotype for the monoclonal antibody REGN10933. Here, we show the convolutional neural network model (Network 6) with optimal performance. Note the several peaks, i.e., escape mutations, at the residues K417, Y453, L455, F486, and Q493 among others. Orange residue index indicates the subset of mutations with additional support as escape mutations from an independent (external) study. The replicated mutations at the residues Y453 and F486 were also the top neural network predictions of escape mutations.

As receptor recognition and antibody escape are closely related viral activities, we implemented Random Forest to calculate the *feature importance* of the 10 convolutional neural network derived mutation-mediated antibody-escape phenotypes as predictors of binding affinity towards the ACE2 receptor. We hypothesized that those Abs that shield the receptor by pre-occupying the spike RBD – ACE2 PD binding interface would weigh more in their contributions to the prediction. Notably, we found that the top 3 Abs (COV2-2499, COV2-2832, and COV2-2050), all entirely targeting the interface based on experimental characterizations^19^, among the neural network derived antibody-escape predictors accounted for nearly 60% of the feature importance score (Figure 6C). Key residues on the spike RBD that are critical binding sites with ACE2 PD include G446, Y449, G496, Q498, and T500 for escape mutations of the top-ranked COV2-2499. Both receptor affinity and the amount of available protein determine the infectivity of SARS-CoV-2 and downstream complications of COVID-19. Notably, among the key binding residues on the spike RBD, these sites of escape mutations for COV2-2499 differed significantly (Mann-Whitney U p=3.2×10^−4^) in their neural network derived binding affinity from the complement set of residues and, even more significantly (p=1.03×10^−19^), in their neural network derived protein expression profile. Thus, the antibody-escape mutations belonged to a sub-cluster within the key binding residue cluster (described above). In addition, we found these residues to be the sites of the peaks for the neural network derived antibody-escape phenotype for COV2-2499, i.e., the predicted escape mutations (Figure 6A). The importance of these residues for viral binding to the host receptor was also supported by results on intermolecular interactions from the molecular dynamics simulations (Figure 5C).

#### Key mutations on ACE2 PD

We comprehensively evaluated mutational effects on the relative binding of ACE2 PD. The corresponding key binding residues in the human ACE2 receptor paired with those of the spike RBD (Figure 5C) differed significantly (Mann-Whitney U p=2.39×10^−7^) in the neural network derived (i.e., estimated, mutation-mediated) relative binding score from the remaining residues, with the key binding residues showing substantially lower variance (i.e., 40% of the variance of the complement set of residues). In addition, we investigated the subset of polymorphic sites in human populations and, using the neural network derived phenotype, identified residues of outlier mutational effects on relative binding score (Figure 5D and 5E). While linear regression (Spearman correlation *ρ* = −0.014) and the multilayer perceptron (Spearman correlation *ρ* = −0.044) failed to make accurate predictions, the best performance achieved from sequence convolutional neural networks was 0.37 (Table S2). The lower phenotypic variance explained by the neural network models on ACE2 PD binding implies a complex sequence-function relationship, with possible contributions from sequence mutations and other (environmental) determinants. Notably, the empirical distribution of the score of the spike RBD and the ACE2 PD (Figure S1) is consistent with the spike RBD displaying more non-conserved mutations than the ACE2 PD, which fits our intuitive understanding of the properties of the virus (which must undergo adaptation to transmission and replication in the host) and the membrane receptor (with its conserved function).

### Independent confirmation of neural network findings

We performed further confirmation of specific neural network predictions and our overall methodology by analyzing an external dataset on an anti-SARS-CoV-2 spike monoclonal antibody REGN10933, which targets the spike-loop region near the edge of the ACE2 interface^29^. We trained neural networks in DMS antibody-escape data^30^ on REGN10933. A map of the neural network derived antibody-escape phenotype for REGN10933 identified several peaks, i.e., escape mutations, at K417, Y453, L455, F486, and Q493 among others. Notably, we found substantial confirmation of the escape mutations at these specific residues in an independent experimental study which had searched for escape mutations through deep sequencing of the passaged virus^31^ (Figure 6D). The replicated mutations at the residues Y453 and F486 were also the top escape mutation predictions.

## Discussion

In this work, we leveraged neural networks to model the biochemical phenotypes of the spike RBD and the ACE2 PD. Mutations can directly influence protein conformational dynamics, stability, and associations/dissociations, i.e., the sequence-structure-phenotype relationship^28^. We examined the prediction performance of several neural network architectures designed to learn the features of the sequence-function landscape. Convolutional neural networks demonstrated superior performance over linear regression or multilayer perceptron in “decoding” the mutational information from sequence to phenotype. Furthermore, the sequence convolutional neural network, in general, outperformed the graph convolutional neural network (Tables S1 and S2). The performance of the graph convolutional neural network might be related to the graph model architecture as well as the physicochemical nature of the predicted phenotype, i.e., the node average model does not sufficiently capture relevant features from a structural perspective for binding affinity. Capturing non-local motifs or long-range dependencies such as allosteric effects in the structural information warrants further study.

Our deep learning experiments confirmed the quality of the AAindex, which captures the intrinsic properties of each amino aside that are independent of the particular sequence, structure, or DMS effect scores. Our shuffling experiments showed that the AAindex significantly improved the prediction. Alternatively put, the use of a randomly generated AAindex was found to be sub-optimal, with the physicochemical properties encoded in the actual AAindex having a significantly greater impact on phenotype prediction. Nevertheless, we observed strong dependence of the biochemical phenotypes on the input sequence data alone, from which the convolutional neural networks could extract features and perform phenotype prediction reasonably well.

Investigating mutational effects is essential for deriving mechanistic insights and elucidating disease pathways. Molecular modeling tools such as molecular dynamics simulations and docking are frequently used towards these ends, with the clear benefit of obtaining atomic-detail understandings of biomolecular systems^9,28^. However, these tools are typically impeded by system complexity and computational demand, so that only a partial structure could be simulated for a limited number of mutational systems. As structural interactions comprise the central elements of molecular modeling, residues on the binding interface with high binding frequency or affinity are likely to be the focus, while dissociation effects like antibody escape are challenging to model. In fact, it has been reported that the rapid spread of COVID-19 has more to do with asymptomatic and pre-symptomatic transmission than enhanced receptor binding^6^. In addition, allosteric effects are difficult to capture in molecular modeling yet critical to viral infection and evolution. It has been reported that the D614G variant that is located far from the RBD of the spike glycoprotein can facilitate ACE2 infection^32^. Similarly, mutations near the ACE2’s chloride-binding sites, which are located far from the spike interface, may also alter the spike RBD binding^18^. Some mutations could become prevalent as selection facilitates receptor binding. For instance, the N501Y mutation has been reported to be responsible for the rapid spread of the virus in southeastern England (which is noteworthy given the result on Tyrosine mutational effect on receptor affinity; Figure 5F), threatening the efficacies of the existing vaccines^33,34^. Deep learning on DMS, as presented here, is a time-efficient and economical way to model high-dimensional mutational data to gain insights into antibody evasion and therapeutic design (Figure 6).

It is worth noting that the tested neural networks achieved much better prediction performance on the spike RBD than on the ACE2 PD. The difference in prediction performance on the two interacting components of the protein complex could result from difference in the data quality, i.e., the DMS experiments had been performed in two separate studies. However, the difference may also be due to a difference in the intrinsic properties of the virus and the receptor. Specifically, the human ACE2 phenotype may be less dependent on (or determined by) sequence mutation data. This finding was also supported by the fact that linear regression or fully connected networks failed to make good prediction for the ACE2 PD.

## Methods

### Datasets

We performed our deep learning experiments on publicly available spike RBD and ACE2 PD DMS datasets^7,18,19^. Binding affinity and protein expression of the spike RBD as well as the relative binding of ACE2 PD dataset were considered. We also leveraged a deep mutational scan of the RBD to determine how variants from spike RBD responded to antibody binding from 10 human monoclonal antibodies, of which nine target SARS-CoV-2 and one targets SARS-CoV. The original mutational effect scores - “escape fraction” - have been normalized via Box Cox transformation.

Models on the avGFP protein (an independent DMS dataset) were also retrained and tested in the RBD expression dataset^16,22^. For this purpose, the wild-type sequence for avGFP was reduced so that the overall sequence length matched that of the spike RBD. Accordingly, the original DMS dataset for avGFP was modified to only include the variant-score profile of the sliced sequence.

### Protein sequence encoding and structural description

AAindex is a resource of 566 physicochemical properties (e.g., polarizability parameter, residue volume, solvation free energy, and other attributes) for each of the 20 amino acids^35^. The AAindex applied in this work underwent dimensionality reduction (from 566 properties to 19 Principal Components)^16^. The wild-type and mutated protein sequence were encoded using one-hot encoding and the AAindex.

### Network architectures and model training

Here, we extend a supervised learning framework for application to interrogating the biochemical phenotypes relevant to the virulence of SARS-CoV-2^16^. We utilized different neural network architectures including linear regression, multilayer perceptron (fully connected), sequence convolutional neural network, and graph convolutional neural network. These architectures can be viewed as directed computational graphs organized as nodes in a series of hidden layers *h*_(*k*)_

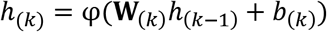

with a matrix of weights **W**_(*k*)_ and biases *b*_(*k*)_. The network architectures are implemented as compositions of functions (equation 1), reflecting the universal approximation properties of the architectures with respect to the compact convergence (compact-open) topology^36^.

Convolutional neural networks perform convolution, activation, pooling, and flattening to learn features from input data. Briefly, convolution is a mathematical operation involving a convolution operator *m* and input *s*:

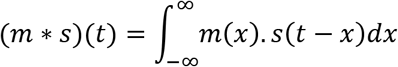

This one-dimensional definition can be extended to arbitrary dimensions. This formulation has a discrete version:

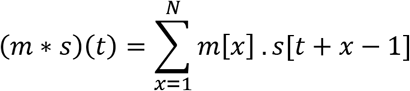

The convolution operation can be viewed as a weight assignment scheme. Different operators or *filters* can be applied to the input data to derive an optimally predictive set of features. Activation introduces non-linearity and pooling helps to reduce overfitting. A fully-connected neural network is then applied to the resulting feature vector (after flattening) to obtain the final prediction.

Minimization of the empirical loss across the training samples

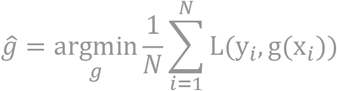

yields an estimate of the mutation- or genetically- determined *ĝ*(*x*), whose properties can be explored (and compared with the original g) and which may enable causal inference on a biochemical phenotype. We trained the models using mean squared error loss and the Adam optimizer with dropout as the regularization method (in particular, dropout rate of 20%). For activation function φ (equation 1), a leaky ReLU was applied, which permits a small, positive gradient for a non-active unit

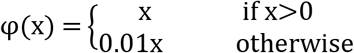

Neural networks were implemented using TensorFlow. We set the maximum possible number of epochs to 400 with early stopping.

The uncertainty associated with the neural network model can be quantified. Let *ω* be the set of all model parameters, i.e., the weights **W**_(*k*)_ and biases *b*_(*k*)_. Then the posterior probability *P*(*ω* | *X*_train_, *Y*_train_) of the set of parameters conditional on the training sequence data *X*_train_ and the training output data *Y*_train_ is, according to Bayes’ rule, given by

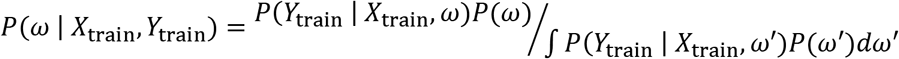

Here, *P*(*ω*) is the *prior* distribution of the set of parameters and *P*(*Y*_train_ | *X*_train_, *ω*) may be assumed to follow

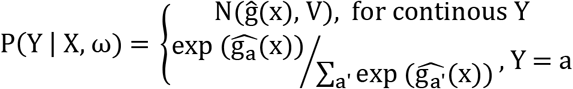

That is, in the continuous case, the likelihood is Gaussian with mean *ĝ*(*x*) and covariance matrix *V* whereas for classification, the conditional probability is a softmax likelihood (assuming the output layer is a softmax layer and the cross-entropy is the loss function). The posterior probability *P*(*ω* | *X*_train_, *Y*_train_) can be calculated using Markov Chain Monte Carlo (MCMC), as the integral (i.e., the so-called *model evidence P*(*Y*_train_ | *X*_train_)) is known to be typically intractable and have no analytical formulation. Prediction of the phenotype *y* for *any* given input *x* can then be quantified through the following inference

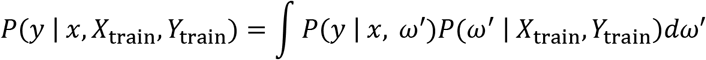

Note the left hand side of this equation can also be sampled using the neural network.

### Joint model consisting of the neural network derived antibody-escape phenotypes

The spike RBD binding affinity was jointly modeled using the convolutional neural network derived (i.e., estimated, mutation-mediated) antibody-escape phenotypes for the ten Abs. The ten “escape fraction” features were scaled and randomly split into 80% training and 20% testing subsets for the Random Forest regressor implemented using sckit-learn^37^. We applied hyper parameter tuning to ensure the best performance.

### AAindex shuffling experiment

We performed a shuffling experiment on the AAindex to assess the extent to which the selected amino acid physicochemical properties impact the performance of the neural network architectures. All elements of the AAindex matrix were re-sampled in-place so that the original covariances among rows or columns were broken. As a result, we generated 100 new (pseudo) AAindex tables. The p-value, i.e., the probability, if the null hypothesis H_0_ (of no gain) is true, that the prediction performance *ρ* (in the test set) is at least as great as the value *ρ*_actual_ from the actual AAindex is calculated as follows

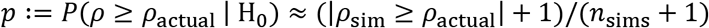

where *ρ*_sim_ is the Spearman correlation from a shuffled dataset, *n*_sims_ is the total number of shuffled datasets, and | | is the count operator.

## Code availability

The code for this project is available from GitHub (https://github.com/gamazonlab/DeepMutScan).

## Author contributions

E.R.G. and B.W. designed the study and wrote the manuscript. B.W. and E.R.G. performed the analyses. E.R.G. supervised and acquired funding for the study.

