## Supplementary Material for "Modeling mutational effects on biochemical phenotypes using convolutional neural networks: application to SARS-CoV-2"

Bo Wang[1] and Eric R. Gamazon[1, 2, 3, 4]

[1]Division of Genetic Medicine, Department of Medicine, Vanderbilt University Medical Center, Nashville, TN, USA

[2]Vanderbit Genetics Institute, Vanderbilt University Medical Center, Nashville, TN, USA

[3]Data Science Institute, Vanderbilt University Medical Center, Nashville, TN

[4]Clare Hall, University of Cambridge, Cambridge, United Kingdom

Send correspondence to:

Eric R. Gamazon <>

Table S1. Neural network performance on spike RBD biochemical phenotypes

| Neural networks | Prediction on binding | Prediction on expression | Architecture |
|---|---|---|---|
| **Linear regression** | 0.488894 | 0.443073 | |
| **Multilayer perceptron** | 0.518265 | 0.49802 | Layer:1 Nodes:100 |
| **Network 1** | 0.731397 | 0.661745 | Layer:1 Kernel:5 Filter:8 |
| **Network 2** | 0.692423 | 0.682139 | Layer:1 Kernel:5 Filter:32 |
| **Network 3** | 0.778903 | 0.704305 | Layer:2 Kernel:5 Filter:32 |
| **Network 4** | 0.717927 | 0.720750 | Layer:3 Kernel:5 Filter:32 |
| **Network 6** | 0.761001 | 0.779276 | Layer:3 Kernel:17 Filter:128 |
| **Network 8** | 0.701028 | 0.678691 | Layer:1 Filter:32 |
| **Network 9** | 0.695742 | 0.680642 | Layer:2 Filter:32 |
| **Network 10** | 0.723501 | 0.838441 | Layer:1 Filter:128 |
| **Network 1-7: sequence convolutional network; Network 8-11: graph convolutional network** | | | |

Table S2. Neural network performance on ACE2 PD binding

| Neural networks | Prediction on binding | Architecture |
|---|---|---|
| **Linear regression** | -0.01453 | |
| **Multilayer perceptron** | 0.044023 | Layer:1 Nodes:100 |
| **Network 5** | 0.371151 | Layer:2 Kernel:17 Filter:128 |
| **Network 6** | 0.343354 | Layer:3 Kernel:17 Filter:128 |
| **Network 7** | 0.328209 | Layer:5 Kernel:17 Filter:128 |
| **Network 2** | 0.207404 | Layer:1 Kernel:5 Filter:32 |
| **Network 3** | 0.310157 | Layer:2 Kernel:5 Filter:32 |
| **Network 8** | 0.090607 | Layer:1 Filter:32 |
| **Network 10** | 0.285816 | Layer:1 Filter:128 |
| **Network 11** | 0.188513 | Layer:2 Filter:128 |
| **Network 1-7: sequence convolutional network; Network 8-11: graph convolutional network** | | |

Table S3. Neural network performance on antibody-escape phenotype for 10 human monoclonal antibodies

| **Antibody types** | **Prediction on antibody escape** |
|---|---|
| **cov2-2050** | 0.246994 |
| **cov2-2082** | 0.09224 |
| **cov2-2094** | 0.315764 |
| **cov2-2096** | 0.19175 |
| **cov2-2165** | 0.123021 |
| **cov2-2479** | 0.130445 |
| **cov2-2499** | 0.260286 |
| **cov2-2677** | 0.29153 |
| **cov2-2832** | 0.125652 |
| **rCR3022** | 0.264203 |
| **Deep Learning experiments performed on network 6** | |

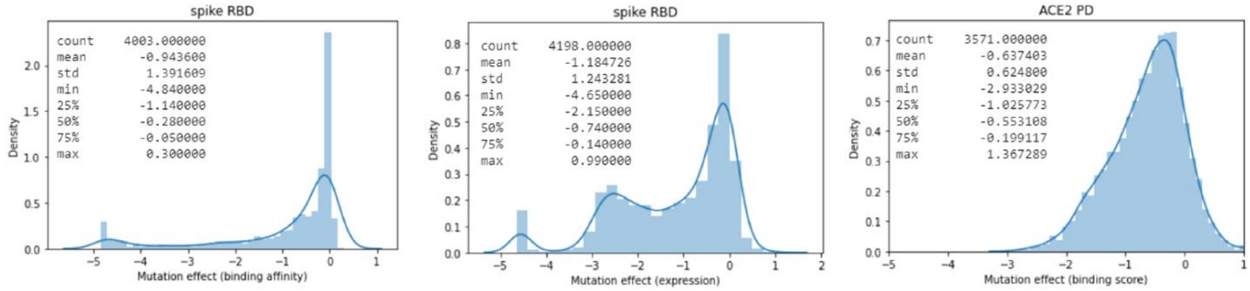


**Figure S1**. Empirical distributions and summary statistics for the biochemical phenotypes of the spike RBD and ACE2.
